# Clinical grade ACE2 as a universal agent to block SARS-CoV-2 variants

**DOI:** 10.1101/2021.09.10.459744

**Authors:** Gerald Wirnsberger, Vanessa Monteil, Brett Eaton, Elena Postnikova, Michael Murphy, Benedict Braunsfeld, Ian Crozier, Franz Kricek, Janine Niederhöfer, Alice Schwarzböck, Helene Breid, Anna Sanchez Jimenez, Agnes Bugajska-Schretter, Alexander Dohnal, Christine Ruf, Romana Gugenberger, Astrid Hagelkruys, Nuria Montserrat, Michael R. Holbrook, Chris Oostenbrink, Robert H. Shoemaker, Ali Mirazimi, Josef M. Penninger

**Author notes:** Correspondence to Gerald Wirnsberger or Josef M. Penninger.

## Abstract

The recent emergence of multiple SARS-CoV-2 variants has caused considerable concern due to reduced vaccine efficacy and escape from neutralizing antibody therapeutics. It is therefore paramount to develop therapeutic strategies that inhibit all known and future SARS-CoV-2 variants. Here we report that all SARS-CoV-2 variants analyzed, including variants of concern (VOC) Alpha, Beta, Gamma, and Delta, exhibit enhanced binding affinity to clinical grade and phase 2 tested recombinant human soluble ACE2 (APN01). Importantly, soluble ACE2 neutralized infection of VeroE6 cells and human lung epithelial cells by multiple VOC strains with markedly enhanced potency when compared to reference SARS-CoV-2 isolates. Effective inhibition of infections with SARS-CoV-2 variants was validated and confirmed in two independent laboratories. These data show that SARS-CoV-2 variants that have emerged around the world, including current VOC and several variants of interest, can be inhibited by soluble ACE2, providing proof of principle of a pan-SARS-CoV-2 therapeutic.

## Introduction

The emergence of SARS-CoV-2 has resulted in an unprecedented COVID-19 pandemic with dire economic, social, and health consequences for hundreds of millions of people. The initial step of SARS-CoV-2 infection is binding of the viral Spike protein to Angiotensin converting enzyme 2 (ACE2) *(1–3)*, followed by proteolytic processing of the trimeric Spike *(4, 5)* and subsequent infection of target cells *(6)*. Inhibition of Spike/ACE2 interaction is the fundamental principle for the activity of neutralizing antibodies induced by all current vaccines *(7)*. Similarly, approved monoclonal antibodies act by blocking the interaction of the cell-entry receptor ACE2 and the viral Spike protein (see https://www.covid19treatmentguidelines.nih.gov/ for further information). Thus, blocking Spike/ACE2 binding has become a central strategy of both vaccine design and multiple therapeutic approaches including ACE2 based therapeutics *(8–12)*. This has created an intense research focus on the molecular details of these processes thereby making the Spike/ACE2 interaction one of the best validated drug targets in biomedicine.

Both vaccines and antibody therapeutics have had an enormous impact and are a remarkable testament to the rapid translatability of basic research. However, although coronaviruses mutate less frequently, as compared to viruses like influenza, many variants of SARS-CoV-2 have emerged throughout the pandemic *(13, 14)* (see also WHO and CDC resources online). Some of these variants have been designated as variants of concern (VOCs) by the WHO because of their increased infectivity and transmissibility. Mutations in the viral Spike protein seem to be of particular relevance in this respect. These mutations do not only affect the infectivity and transmissibility of SARS-CoV-2, but also reduce the potency of vaccines, convalescent sera, and monoclonal antibody therapeutics *(14–20)*. The recent emergence of the Delta variant and symptomatic and sometimes even severe infections of doubly vaccinated people *(21–23)* is one such example and more such variants will likely develop due to the evolutionary pressure of mass vaccinations of hundreds of millions of people during an active pandemic. To prevent further severe disruptions to life and economies due to SARS-CoV-2 infections, it is therefore paramount to design universal strategies for the prevention and treatment of current VOCs and possibly even to variants that will emerge in the future.

Here we report that soluble ACE2 (APN01), already being tested in clinical trials (NCT04335136), binds receptor binding domain (RBD) and full-length Spike proteins of SARS-CoV-2 variants with increased affinity when compared to the SARS-CoV-2 reference strain Spike and effectively neutralizes infections of all tested variants. We also report in our accompanying paper (Shoemaker et al.) that the same clinical grade APN01 can prevent COVID-19 symptoms in a faithful SARS-CoV-2 infection animal model and can be administered as an aerosol directly into the lungs of infected individuals to neutralize SARS-CoV-2. Clinical Phase I testing of this inhalation approach is currently underway. Since Spike/ACE2 interaction is the crucial first step of viral infection, the viral Spike cannot mutate out of ACE2 binding, without a loss in infectivity and tissue tropism. Our data provide the blueprint for a universal anti-COVID-19 agent with the potential to treat or even prevent infections against, in essence, all current and future SARS-CoV-2 variants.

## Results

### Enhanced affinity of clinical grade ACE2 to the Spike RBDs of emerging SARS-CoV-2 variants

Viral evolution of SARS-CoV-2 has been demonstrated to be focused on the Spike protein which is instrumental for the early steps of viral infection *(24, 25)*. Many single or compound mutations, especially in the RBD of the viral Spike, have been described and either hypothesized or demonstrated to affect binding to the cell entry receptor ACE2 (see *(14)* for a review). To systematically test whether these emerged variants affect Spike/ACE2 interactions, we selected viral variants that have been described in the literature and in various databases. The RBD variants analyzed in this study and the location(s) of the respective mutations are depicted schematically (Figure 1a) and also in a 3D model of the viral Spike RBD (Figure 1b). In an initial set of experiments, we performed ELISA analyses with plate-bound ACE2/APN01 to evaluate the impact of the indicated single and compound mutations on Spike RBD/ACE2 interactions. Intriguingly, almost all tested variant RBDs exhibited increased binding to APN01 (Figure 1c). These results prompted us to extend the number of analyzed variants to include current VOCs and to biophysically characterize RBD/ACE2 interactions using Biacore Surface Plasmon resonance (SPR) analysis.

**Figure 1.**
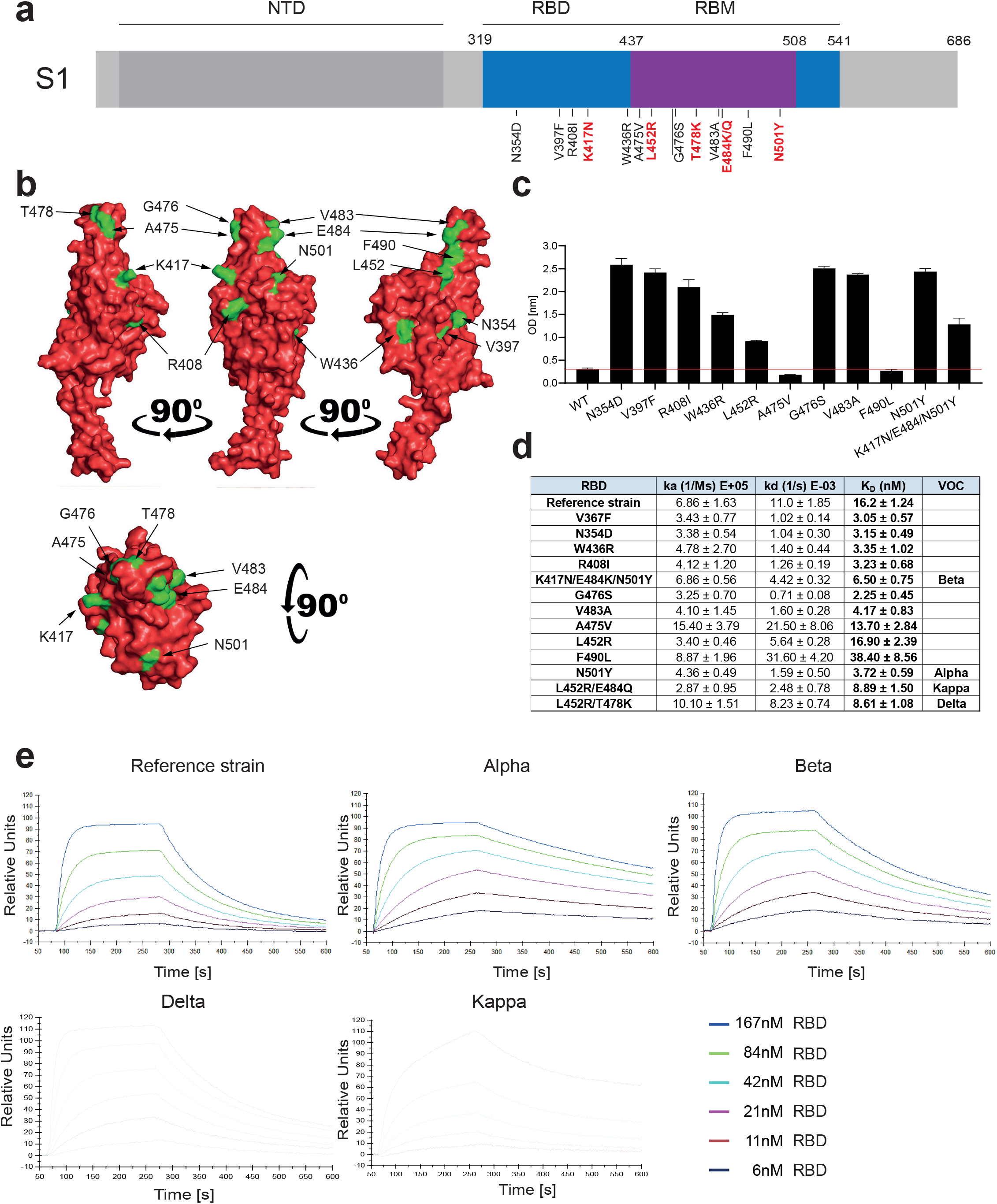
Increased affinity of APN01 interactions with SARS-CoV-2-RBD variants. **(a)** Schematic depicts structure of the SARS-CoV-2 spike protein S1 domain. Indicated is the amino terminal domain (NTD), the receptor binding domain (RBD) in blue and within the RBD the receptor binding motif (RBM) in purple. Numbers above depict domain boundaries. Mutations within the RBD/RBM are indicated below with observed amino acid exchanges. Shown in red are mutations observed in Variants of Concern (VOC). **(b)** PyMOL rendered visualization of the SARS-CoV-2 RBD. Rendering depicts the SARS-CoV-2 RBD with mutation sites shown in green. **(c)** ELISA analysis showing the binding strength of SARS-CoV-2 RBD carrying the indicated mutations to APN01. Axis labels indicate the SARS-CoV-2 RBD variant substitutions tested. **(d)** Surface Plasmon Resonance analysis to derive kinetic constants (k_a_, K_d_) and affinity values (K_D_) of SARS-CoV-2 RBD/APN01 interaction. The table lists both the tested variants and the introduced amino acid substitution as well as the designation of the respective Variants of Concern mutations tested in this study. Reference strain RBD sequence corresponds to the Wuhan SARS-CoV-2 isolate (e) Representative SPR sensorgram images for the SARS-CoV-2 RBD/APN01 interaction.

For comparative kinetic binding analysis of SARS-CoV-2 RBD variants, dimeric APN01 was covalently coupled as ligand to optical sensor chips. Commercially available or in house purified RBDs containing the amino acid changes described in Figure 1a and 1b were passed as analytes over the immobilized APN01 ligand in twofold serial dilutions. These proteins contained previously identified RBD mutations of current VOCs, as well as amino acid substitutions identified in variants of interest (VOI). Binding on-rates (association constants; k_a_), off-rates (dissociation constants; k_d_) and binding affinities (K_D_ reported as nM) of SARS-CoV-2 RBD/ACE2 interactions were determined by mathematical sensorgram fitting, applying a monomeric Langmuir 1:1 interaction model (A + B = AB) using BiaEvaluation 4.1 software. The results are summarized in Figure 1d, listing both the tested variants and the introduced amino acid substitution as well as the designation of the respective clinical VOC isolates that were tested with SPR analysis. Of note, the reference RBD sequence corresponds to the original Wuhan SARS-CoV-2 isolate. Importantly, our SPR analyses showed that affinities of all tested VOC-RBDs to APN01 are substantially increased, with the Alpha variant RBD showing the highest affinity (Figure 1d,e). Of note, among these VOC RBDs we observed a significantly lower off-rate for the Alpha variant and a significantly faster on-rate for the Delta variant. Combined with the increased binding affinity, changes in these kinetic parameters might contribute to the enhanced infectivity of VOC. These data show that, as expected and also in part reported by others *(26–30)*, viral evolution of the Spike protein led to an increase in Spike RBD/ACE2 binding affinity as well as altered kinetic constants of RBD/APN01 interaction, especially observed for VOC.

### Enhanced binding of SARS-CoV-2 VOC Spike trimers to clinical grade ACE2/APN01

SARS-CoV-2 RBD binding to ACE2 occurs in the context of trimeric Spike proteins in the pre-fusion conformation. Spike trimer structures have been solved in receptor bound or unbound forms *(31, 32)* and it has been demonstrated that the conformation of the Spike (open or closed) is of critical relevance for receptor interactions *(33, 34)*. Importantly, mutations outside the RBD that alter the Spike conformation have been reported to also affect viral infectivity and receptor binding *(35, 36)*. To test whether our observations on the increased affinity of VOC RBD/APN01 interaction is also observed in the context of the full-length Spike, we assessed APN01 binding to recombinant prefusion trimeric SARS-CoV-2 Spike proteins. Since APN01 is a dimeric molecule thus allowing for bivalent target interaction, the VOC trimeric pre-fusion Spike variant proteins were immobilized to an optical sensor chip surface by covalent amine coupling. APN01 was passed over the immobilized Spike proteins in serial dilution in single binding cycles. Using BiaEvaluation 4.1 software, subsequent kinetic analysis was carried out by sensorgram fitting applying a Langmuir binding and a bivalent analyte binding model. Kinetic binding constants derived from the Langmuir model represent the apparent affinity. The bivalent analyte model calculates separate kinetic constants for the affinity determining first step A + B = AB and the avidity determining second step AB + B = AB_2_ of the binding process.

Structural rendering of the trimeric full-length SARS-CoV-2 Spike protein and the positions mutated in the various strains of SARS-CoV-2 used in this study are shown in Figure 2a. Lineages, mutations, and sources of the SARS-CoV-2 isolates including VOCs and VOIs used in our biophysical analyses are listed in Figure 2b. SPR analysis showed strong binding of APN01 to all tested variants of the pre-fusion Spike trimers including the original Wuhan viral isolate trimer as well as the Alpha, Beta, Gamma, Delta, and Kappa variants (Figure 2c). Sensorgram fitting showed enhanced apparent affinity (Langmuir model) and avidity (bivalent analyte model) of all VOC trimeric Spike proteins, except for the Kappa VOI (Figure 2c, d). These results demonstrate that, when compared to Spike trimers of the reference strain, dimeric recombinant soluble human ACE2 (APN01) binds to the pre-fusion Spike trimers from all current VOCs with increased affinity/avidity.

**Figure 2.**
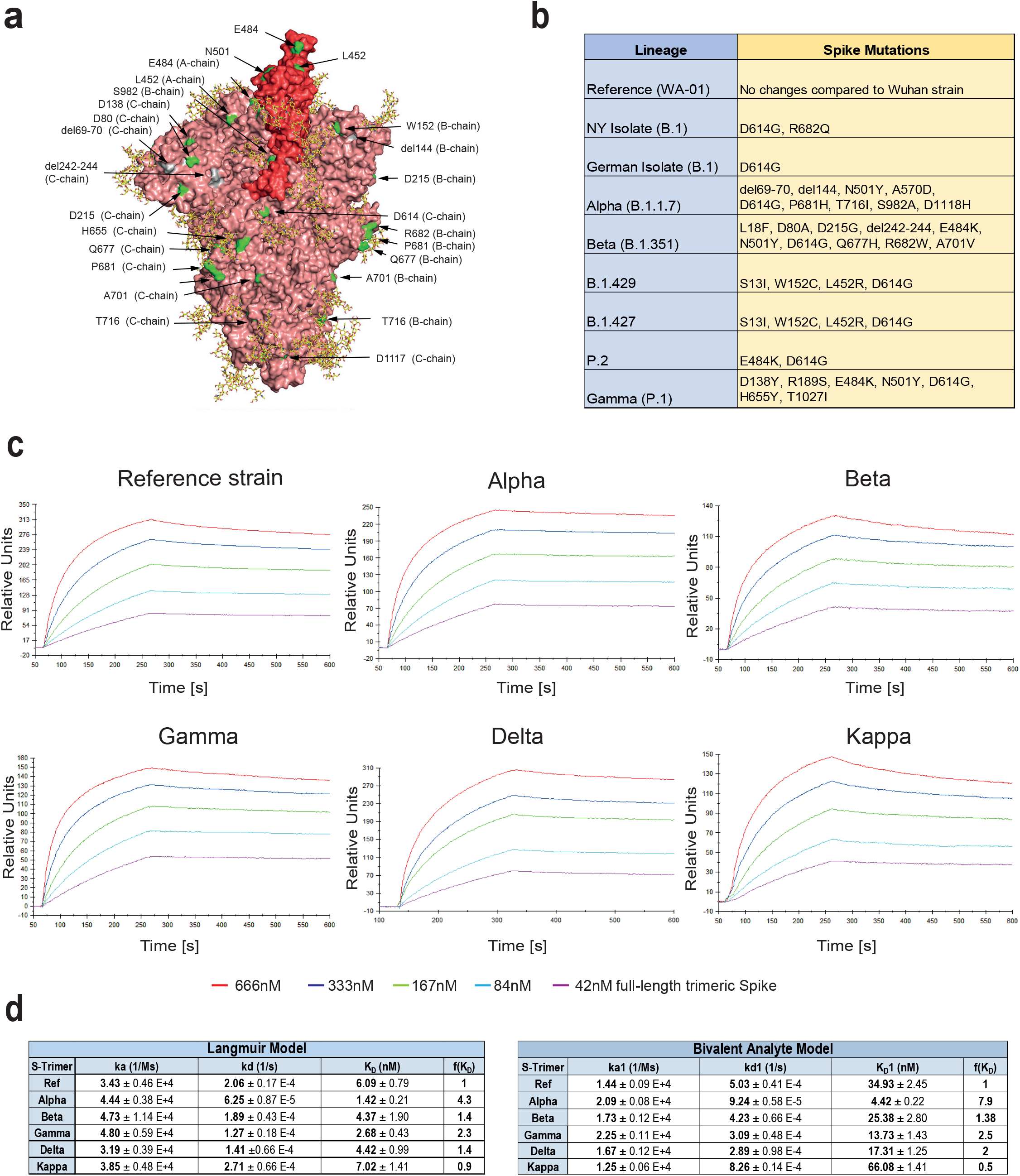
Increased binding affinity of APN01 to full-length pre-fusion trimeric Spike proteins from SARS-CoV-2 variants of concern. **(a)** PyMOL rendering of the trimeric full-length SARS-CoV-2 Spike protein. One RBD is shown in red. Indicated in green are positions mutated in the various strains of SARS-CoV-2 used in experiments in this study. Depicted in yellow are the glycan-modifications of the spike protein. **(b)** Table lists the SARS-CoV-2 strains used in this study **(c)** Representative sensorgram images for the SPR analysis conducted with full-length trimeric spike proteins in pre-fusion state with APN01. Reference strain corresponds to original Wuhan viral isolate spike sequence. Indicated are VOC Alpha, Beta, Gamma, and Delta, as well as the variant Kappa. **(d)** Tables listing k_a_, k_d_, as well as k_D_ values for the interaction of APN01 and full-length trimeric spike proteins. Values are derived from calculations based upon the Langmuir (upper table) or Bivalent Analyte sensorgram fitting (lower table).

### ACE2/APN01 effectively neutralizes SARS-CoV-2 VOCs

We have previously reported that clinical grade APN01 can effectively reduce the SARS-CoV-2 viral load in VeroE6 cells and 3D organoids in a dose dependent manner, using a reference virus isolated early during the pandemic *(37)*. This virus carries the same Spike sequence as the originally reported virus isolated in Wuhan. To test whether APN01 can also neutralize variant clinical SARS-CoV-2 isolates including VOC strains (see Figure 3a for a list of tested strains), we performed neutralization assays in VeroE6 cells and compared its inhibitory potency side by side to our reference strain. APN01 indeed potently neutralized all the SARS-CoV-2 isolates we tested in VeroE6 cells (Figure 3b). Intriguingly, this inhibition was markedly enhanced against all the VOCs tested with sometimes up to 20 times lower IC_50_ and IC_90_ values (Figure 3c).

**Figure 3.**
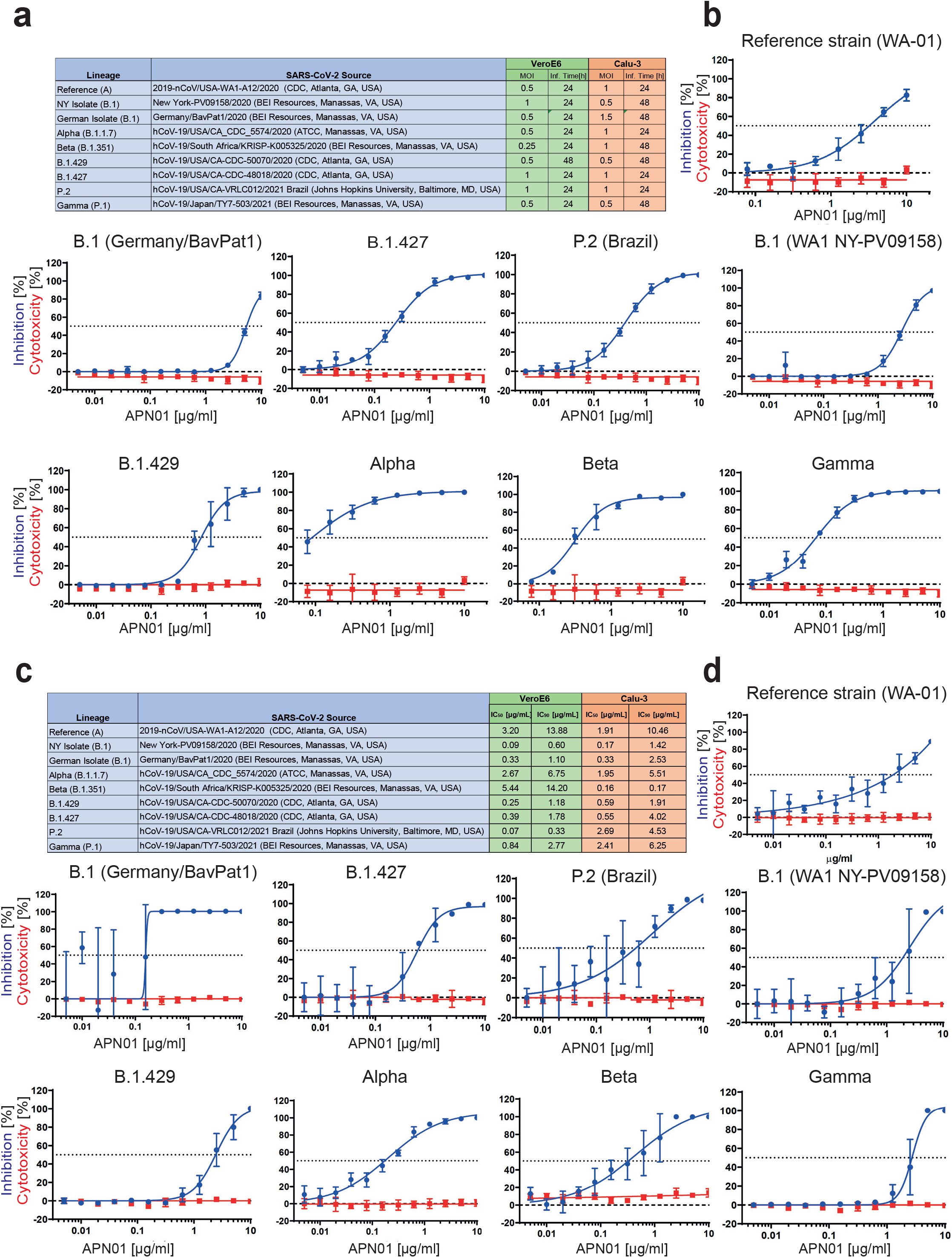
Increased neutralisation potency of APN01 towards SARS-CoV-2 variants. **(a)** Table depicts source of the tested viral isolates, as well as multiplicity of infection (MOI) and the infection time used in these experiments for both VeroE6 and Calu-3 cells. **(b)** Panels depict both neutralization of the indicated SARS-CoV-2 isolates (blue line) as well as cytotoxicity of APN01 (red line) in VeroE6 cells. Analysis was done in quadruplicate with mean and standard deviations shown. Y-axis depicts the percentage of neutralization and cytotoxicity, respectively. **(c)** Table depicts IC_50_ and IC_90_ values for APN01 mediated neutralization of viral infection in VeroE6 and Calu-3 cells. **(d)** Same as (a) but conducted with epithelial lung cancer cell line Calu-3.

To extend our results to a physiologically more relevant cell system, we infected Calu3 human lung epithelial cells with the reference SARS-CoV-2 isolate and the indicated SARS-CoV-2 variants. We again observed that clinical grade soluble APN01 potently reduced viral load of all tested variants in a dose dependent manner (Figure 3c, d). Importantly, the observed neutralization potency closely correlated with the Spike/APN01 binding affinity as assessed via SPR. In some cases, we detected 10 to 20 times lower IC_50_ and IC_90_ values when compared to values obtained with the SARS-CoV-2 reference strain. These data show that ACE2/APN01 not only binds significantly stronger to RBD or full-length Spike proteins of the tested variants, but also more potently inhibits viral infection by these strains in both VeroE6 and a human lung epithelial cells.

### Independent validation studies

Experimental set-ups and model systems as well as culture conditions sometimes have a dramatic impact on experimental results. To ensure the reproducibility of our results we conducted confirmatory experiments in a different and independent laboratory. These validation experiments were performed at the Karolinska Institutet, Stockholm, where VeroE6 cells were infected with reference virus isolated from the first Swedish patient. This virus was previously reported *(37, 38)* and carries the same Spike amino acid sequence as reported for the first Wuhan virus isolate. Experiments were performed at different Multiplicities of Infection (MOIs) to test the inhibitory potency of APN01. As reported before *(37)*, APN01 markedly reduced viral replication of the SARS-CoV-2 reference strain in a dose dependent manner (Figure 4a,b). Importantly, the inhibitory potency of APN01 was again significantly increased towards the VOC Alpha and Beta, providing an independent validation of the neutralization results. This increase in neutralization potency of APN01, when the reference virus was compared to the VOC strains Alpha and Beta, could be observed at all APN01 concentrations and MOIs tested (Figure 4a, b, c). These results, using different viral isolates as well as different experimental procedures, independently confirm that clinical grade soluble human APN01 strongly blocks SARS-CoV-2 infections of recently emerged VOC.

**Figure 4.**
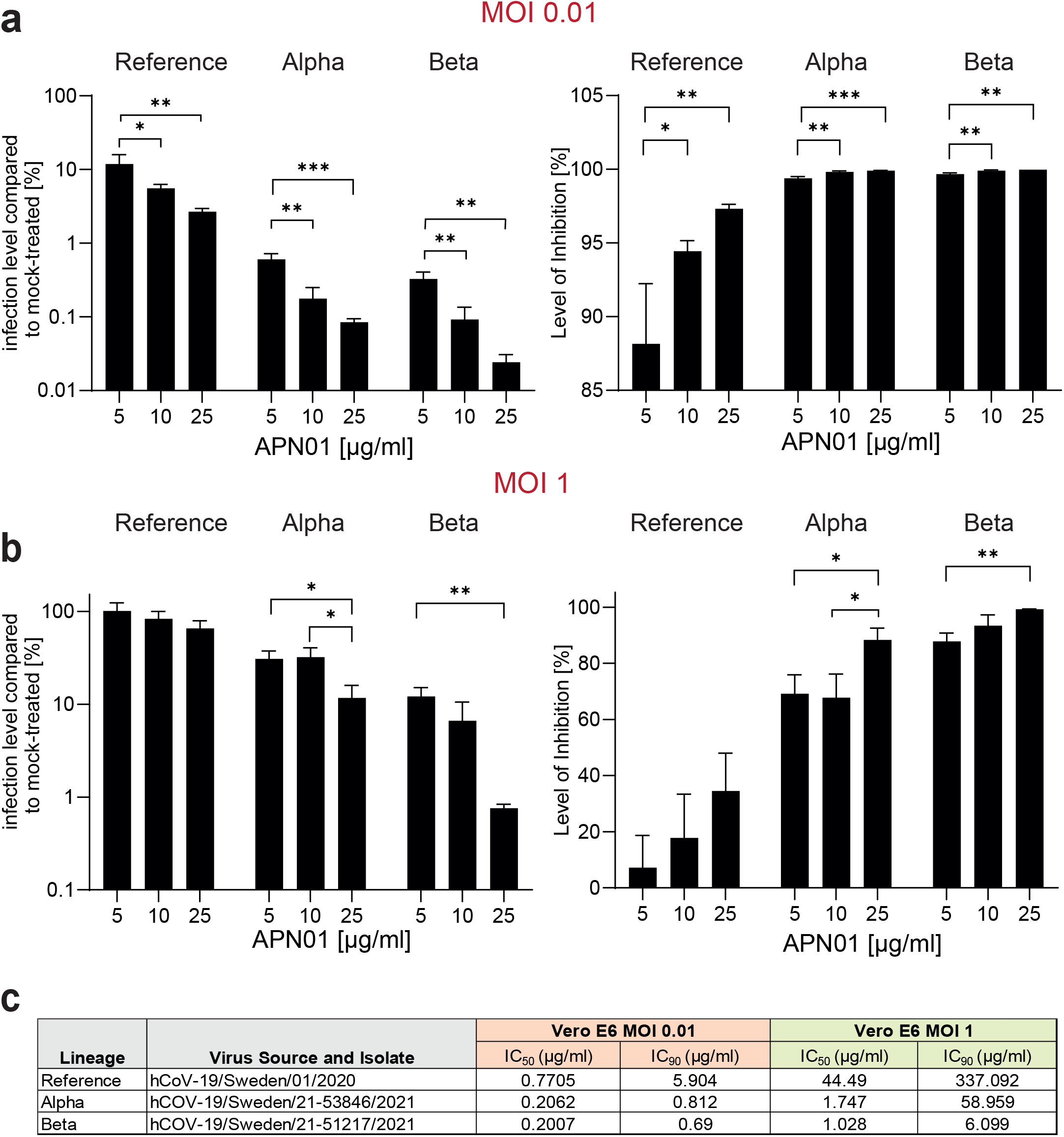
Increased potency of APN01 against SARS-CoV-2 VOCs. **(a,b)** Diagrams depict the level of infection with the indicated SARS-CoV-2 isolates at MOI 0.01 (a) and MOI 1 (b) of VeroE6 cells in the presence of increasing concentrations of APN01 as compared to infections in the absence of APN01. Shown are means of triplicate analyses with standard deviations. Statistical significance is indicated by asterisks (p-value < 0.05: *; p-value < 0.001: *** as calculated with one-way ANOVA) **(c)** List and source of strains used at the Karolinska Institutet and IC_50_ and IC_90_ values obtained for the indicated MOIs. See Materials and Methods section for a detailed list of viral mutations for the strains used.

## Discussion

Since emerging from Wuhan, China, in December of 2019, SARS-CoV-2 has been causing devastating severe respiratory infections in humans worldwide. Multiple variants of SARS-CoV-2 have emerged and circulate around the world throughout the COVID-19 pandemic with some strains displaying even greater infectivity and transmissibility (see https://www.cdc.gov/coronavirus/2019-ncov/variants/variant-info.html and https://www.who.int/en/activities/tracking-SARS-CoV-2-variants/ for further information). Some of these variants have been classified by the WHO as *Variants of Concern* (VOC), defined as “*a variant for which there is evidence of an increase in transmissibility, more severe disease, significant reduction in neutralization by antibodies generated during previous infection or vaccination, reduced effectiveness of treatments or vaccines, or diagnostic detection failures***”** or *Variants of Interest* (VOI), defined as **“***a variant with specific genetic markers that have been associated with changes to receptor binding, reduced neutralization by antibodies generated against previous infection or vaccination, reduced efficacy of treatments, potential diagnostic impact, or predicted increase in transmissibility or disease severity*.” The current VOCs are B.1.1.7 (Alpha), B.1.351 (Beta), P.1 (Gamma) and B.1.617.2 (Delta), whereas multiple VOI are currently circulating including B.1.526 (Iota), B.1.427, B.1.429, B.1.617.1 (Kappa), B.1.617.3, B.1.525 (Eta), B.1.621 (Mu), or C.37 (Lambda). Both large-scale vaccination programs as well as emerging treatments will presumably drive further molecular evolution of SARS-CoV-2. It is therefore to be expected that novel variants will arise, some of which will rapidly spread and even re-infect fully vaccinated people, sometimes leading to severe breakthrough COVID-19, as is currently seen with the Delta variant *(21, 22)*. In this arena of global vaccination efforts and antibody therapeutics in multiple permutations, it therefore remains critical to identify and design universal strategies that might help prevent and treat infections with all current and potential future variants.

The SARS-CoV-2 Spike protein interacts with high affinity with its main entry receptor ACE2, followed by a subsequent membrane fusion step. Although alternative ACE2 independent modes of viral uptake and infection have been reported, genetic modeling in ACE2 deficient mice has provided unambiguous evidence that ACE2 is the essential receptor for SARS-CoV-2 infections and subsequent development of COVID-19 in mice (Gawish et al. 2021, BioaRxiv.org). Indeed, most neutralizing antibodies from vaccinations, convalescent plasma therapies, and monoclonal antibodies or nanobodies interfere with the Spike/ACE2 interaction *(14)* and have, due to their therapeutic or prophylactic efficacy, provided further evidence for the essential role of ACE2 as the relevant *in vivo* receptor for SARS-CoV-2 and COVID-19 in humans. Conceptually, all SARS-CoV-2 variants and “escape mutants” still bind to ACE2 *(39–46)*, in part already validated in various studies *(41, 46, 47)*. ACE2 is a carboxypeptidase that controls angiotensin II peptide levels and thereby is involved in critical aspects of physiology such as blood pressure control or sodium retention as well as disease processes including heart failure, blood vessel and kidney pathologies in diabetes, tissue fibrosis, or regulation of inflammatory cytokines *(48)*. Our group was the first to show that soluble dimeric ACE2 protects mice from acute lung injury and Acute Respiratory Distress Syndrome (ARDS) *(49)* which triggered preclinical and clinical development of recombinant human soluble ACE2 *(50, 51)*, termed APN01, for lung disease. It was therefore critical to systematically assess whether APN01 could indeed bind to the RBD and more importantly full-length Spike of emerged SARS-CoV-2 variants and neutralize infections with these variants – especially recently emerged VOC. Our data show that APN01 associates with the RBD and prefusion trimeric Spike of all variants tested, often at markedly increased affinity and avidity, which is in line with the enhanced infectivity of these variants. Importantly, using two different cell types including lung epithelial cells, APN01 showed markedly improved efficacy to block infection of all VOC, demonstrating that the prediction holds true – clinical grade APN01 can effectively block all tested SARS-CoV-2 variants and this inhibition is markedly improved against all tested VOCs and VOIs.

APN01 has now undergone phase 2 testing in severe (WHO scores 4, 5, and 6) COVID-19 patients using intravenous infusions, a study that is now being expanded to a larger patient population. In our accompanying paper we have developed a new formulation of APN01 that can be inhaled as an aerosol to directly interfere with the earliest steps of viral infection and prevent lung damage and spreading to other organ systems. Moreover, our accompanying paper demonstrates preclinical efficacy of APN01 lung administration in a mouse adapted severe COVID-19 model, thereby demonstrating proof of concept for phase 1 clinical inhalation trials in humans that are currently ongoing. Of note, the FDA recently expanded emergency use authorization for a two-antibody “cocktail” to allow its use in patients seeking protection from COVID-19 following exposure to someone infected with SARS-CoV-2, providing a framework for preventive and post-exposure prophylactic strategies. However, structural and functional analyses of the VOCs reveal decreased efficacy and viral escape from therapeutic antibodies or antibodies generated from natural infection and vaccinations. Further viral evolution, especially under selective pressure by both widespread vaccination programs and monoclonal antibodies or antibody cocktails is likely to drive the emergence of resistant novel variants and strains. In particular, the Delta VOC strain has already substantially contributed to a new global wave of infection and concerningly large numbers of re-infections. Our data show that clinical grade soluble ACE2/APN01 blocks infectivity of all tested variants supporting the notion that this therapeutic is inherently resistant to escape mutations that constitute a major problem for both vaccines and antibody-based therapeutics. Data provided in this study and in our accompanying manuscript as well as clinical data already generated for APN01 set the stage for the development of universal and pan-variant SARS-CoV-2 prevention and therapy.

### Limitations of this study

Our study used two different cell types and should be expanded using additional human cell types and also organoids. Moreover, to indeed make a claim on universality, additional variants should (and can) be tested using affinity/avidity measurements and neutralization assays.

## Acknowledgements

We like to thank employees of Apeiron Biologics’ R&D department for input and critical discussion during the course of this study. We would also like to acknowledge Mr. Gregory Kocher (IRF-NIAID-NIH) for growing SARS-CoV-2 virus variants.

## Funding

J.M.P. and A.H. and the research leading to these results has received funding from the T. von Zastrow foundation, the FWF Wittgenstein award (Z 271-B19), the Austrian Academy of Sciences, the Innovative Medicines Initiative 2 Joint Undertaking (JU) under grant agreement No 101005026, and the Canada 150 Research Chairs Program F18-01336 as well as the Canadian Institutes of Health Research COVID-19 grants F20-02343 and F20-02015. Additionally, this project has received funding from the Innovative Medicines Initiative 2 Joint Undertaking (JU) under grant agreement no. 101005026. The JU receives support from the European Union’s Horizon 2020 research and innovation program and EFPIA. Parts of this project have been funded with federal funds from the National Cancer Institute, National Institutes of Health, under Contract No. 75N91019D00024, Task Order No. 75N91019F00130. The content of this publication does not necessarily reflect the views or policies of the Department of Health and Human Services, nor does mention of trade names, commercial products, or organizations imply endorsement by the U.S. Government (IC).

## Author contributions

G.W. developed the concept for this study, designed and analyzed experiments, and wrote the manuscript with input from all authors. J.N. performed experiments and analyzed data. A.S. performed experiments and analyzed data. H.B. performed experiments and analyzed data. R.G. provided conceptual input to this study. A.B.-S. and A.D. provided conceptual input to this study. A.S.J. performed experiments and analyzed data. J.P.M. provided conceptual input and wrote the manuscript with input from all authors. A.M. designed experiments, provided conceptual input, and analyzed data. V.M. planned, performed, and analyzed data. M.H. designed and analyzed experiments and provided conceptual input. I.C. designed and analyzed experiments and provided conceptual input. E.P. designed, performed, and analyzed experiments. C.O. and B.B. designed and performed structural modeling work. F.K. and C.R. designed, performed, and analyzed biophysical experiments and provided conceptual input. N.M. provided conceptual input to this study.

## Competing interests

J.M.P. declares a conflict of interest as a founder and shareholder of Apeiron Biologics. G.W., R.G., A.S., J.N, A.S.J, and H.B. are employees of Apeiron Biologics. Apeiron holds a patent on the use of ACE2 for the treatment of various diseases and is currently testing APN01 (soluble recombinant human ACE2) for treatment in COVID-19. The other authors do not declare any conflict of interest.

## Materials and Methods

### Surface Plasmon Resonance Analysis

Recombinant SARS-CoV-2 spike proteins (His-tagged) were purchased from Acro Biosystems Inc. (Newark, USA) or produced inhouse by overexpression of respective constructs in HEK293T cells followed by purification via nickel NTA chromatography and size exclusion chromatography. The purity of recombinant proteins was documented by SDS Page analysis. SPR measurements were performed on a Biacore 3000 instrument (GE Healthcare). For comparative kinetic analysis of SARS-CoV-2 RBD variants, APN01, which does not contain a capture tag was immobilized on optical sensor chip surfaces by covalent coupling at pH = 4.5 at ligand densities of 2672 ± 145 RU following the Biacore amine coupling protocol. Amine activated flow cell 1 (FC1) was used as a reference to allow for the generation of background-subtracted binding sensorgrams. SARS-CoV-2 RBD protein variants were passed over the immobilized APN01 ligand as analytes in twofold serial dilution (167 nM, 84 nM, 42 nM, 21 nM, 11 nM, and 6nM) in single binding cycles, at a flow rate of 30 μl/minute in HBS-EP buffer (0.1 M HEPES, 1.5 M NaCl, 0.03 M EDTA and 0.5% v/v Surfactant P20). Bound analyte was removed after each cycle by surface regeneration with 3M MgCl_2_. Reference flow cell (FC1) subtracted sensorgram overlays with additional correction by subtracting buffer (c=0) sensorgrams (double referencing) were generated and used for kinetic binding analysis. Subtraction spikes occurring at the injection start were removed in the sensorgrams shown in the figures. Kinetic binding constants (k_a_, k_d_ and K_D_) were generated by mathematical sensorgram fitting. Generally, a Langmuir 1:1 interaction model (A + B = AB) was applied, using BiaEvaluation 4.1 software. A series of 4 curve fittings was performed for each binding reaction, using a simultaneous single fitting algorithm for each of the sensorgram overlays. Mean values and standard deviations were obtained from fitting runs with Chi^2^ values ≤ 3% of R_max_. Binding affinities (reported as nM) were calculated from on- and off-rate constants.

SPR analysis of APN01 binding to recombinant full-length trimeric SARS-CoV-2 spike proteins was performed with the spike proteins immobilized to CM5 optical sensor chip surfaces by covalent amine coupling to reach a surface density of 900 – 1000 RU. APN01 was passed over the immobilized spike proteins as dimeric bivalent analyte at five concentrations (42 nM, 84 nM, 167 nM, 333 nM and 666 nM) in repetitive single binding cycles. Using BiaEvaluation 4.1 software, kinetic analysis was carried out by applying a Langmuir 1:1 (apparent affinity) and a bivalent analyte binding algorithm, which separates first step binding (A + B = AB; affinity) from the second step binding (AB + B = AB_2_; avidity).

### Cell lines and cell culture

Infection and APN01 mediated viral neutralisation assays were conducted at the Integrated Research Facility at Fort Detrick (IRF-Frederick) of the National Institute of Allergy and Infectious Diseases (NIAID) or at the Department of Laboratory Medicine (Unit of Clinical Microbiology) of the Karolinska Institutet and Karolinska University Hospital. Vero E6 cells were cultured in DMEM medium (Gibco, Gaithersburg, MD, USA or Thermofisher) containing 10% fetal bovine serum (FBS). Calu-3 cells (HTB-55; American Type Culture Collection [ATCC], Manassas, VA, USA) were cultured in DMEM F12 Medium (ATCC) with 20% FBS.

### Viral neutralization experiments conducted at the Karolinska Institutet

24h after seeding of Vero E6 cells (5×10^4^ per 48 well) APN01 was mixed with viral particles of the indicated strains at the given concentrations in DMEM Medium (Thermofisher) containing 5% FBS in 100μl per well and incubated for 30min at 37°C. After the incubation period medium was removed from Vero E6 cells, cells were washed once with PBS to remove any non-attached cells and virus/APN01 mixtures were added to the cells. Cells were incubated with virus for 15h, after which cells were washed 3 times with PBS and lysed the Trizol, subsequently. RNA was extracted using the direct-zol RNA kit (Zymo Research) and assayed by qRT-PCR as previously described (Monteil et al, Cell, 2020). Half-maximal inhibitory concentration (IC_50_) and inhibitory concentration 90 (IC_90_) were calculated using GraphPad Prism Software (La Jolla, CA).

### Viral neutralization experiments conducted at the IRF-Fredrick

For infection experiments Vero E6 cells were seeded at 6,000 cells in 30μl in 384 wells 24h prior to infection and Calu-3 cells were plated at 10,000 cells per well in 30μl 48h prior to infection in their respective culture media. For APN01 neutralization experiments APN01 was diluted 2-fold in an eight- or twelve-point dose curve. Each condition was tested in quadruplicate (n=4) with an efficacy plate and a mock infected cytotoxicity plate run in parallel. Strains for infection and MOIs are listed in a table in the respective figures. Suitable MOIs were optimized previously for each cell line and virus using serial dilutions and staining for SARS-CoV-2 nucleoprotein, as described below. The viral inoculum was diluted in the respective cell culture medium containing indicated doses of APN01 and pre-incubated for 60min. 20μl of the virus/APN01 mixture was transferred directly to plates containing cells to reach a final volume of 50μl. After infection cells were incubated for 24h or 48h, in accordance with optimal virus and cell-line infection conditions. Cells were then fixed in 10% formalin and stained with a SARS-CoV-1 nucleoprotein-specific antibody (cross reactive to SARS-CoV-2; see antibody list for further information), followed by a secondary antibody conjugated with fluorophore and/or horseradish peroxidase (HRP). Fluorescence was quantitated using a PerkinElmer Operetta high-content imaging system (PerkinElmer, Massachusetts, USA). Chemiluminescence was read on a Tecan M1000 plate reader (Tecan, Switzerland). Cytotoxicity on mock infected plates was determined using the Promega CellTiter-Glow Luminescent Cell Viability Assay (Promega, Madison, Wisconsin, USA). Half-maximal inhibitory concentration (IC_50_) and 50% cytotoxic concentration (CC_50_) were calculated as described by Covés-Datson et al., 2019, using GraphPad Prism Software (La Jolla, CA). Z’ factor scores were assessed as quality-control parameters for each plate of each run. All plates included in the report passed quality-control criteria.

### Viral Strains and isolates

Sources and Strains used in the experiments at the NIAID and Karolinska Institutet are indicated in the respective figures, including a list of SARS-CoV-2 spike mutations identified by sequencing of viral isolates (NIAID). Mutations in isolates used at the Karolinska are as follows (with Spike mutations indicated in bold):

| Position | Wuhan | Alpha Variant | Type | Gene |
| --- | --- | --- | --- | --- |
| 241 | C | T | SNP | ORF1ab |
| 913 | C | T | SNP | ORF1ab |
| 3037 | C | T | SNP | ORF1ab |
| 3267 | C | T | SNP | ORF1ab |
| 3743 | C | T | SNP | ORF1ab |
| 5388 | C | A | SNP | ORF1ab |
| 5986 | C | T | SNP | ORF1ab |
| 6954 | T | C | SNP | ORF1ab |
| 11287 | GTCTGGTTTT | G | DEL | ORF1ab |
| 14408 | C | T | SNP | ORF1ab |
| 14676 | C | T | SNP | ORF1ab |
| 15279 | C | T | SNP | ORF1ab |
| 16176 | T | C | SNP | ORF1ab |
| **21764** | **ATACATG** | **A** | **DEL** | **S** |
| **21990** | **TTTA** | **T** | **DEL** | **S** |
| **23063** | **A** | **T** | **SNP** | **S** |
| **23271** | **C** | **A** | **SNP** | **S** |
| **23403** | **A** | **G** | **SNP** | **S** |
| **23604** | **C** | **A** | **SNP** | **S** |
| **23709** | **C** | **T** | **SNP** | **S** |
| **24506** | **T** | **G** | **SNP** | **S** |
| **24914** | **G** | **C** | **SNP** | **S** |
| 27972 | C | T | SNP | ORF8 |
| 28048 | G | T | SNP | ORF8 |
| 28111 | A | G | SNP | ORF8 |
| 28280 | GAT | CTA | MNP | N |
| 28881 | GGG | AAC | MNP | N |
| 28977 | C | T | SNP | N |

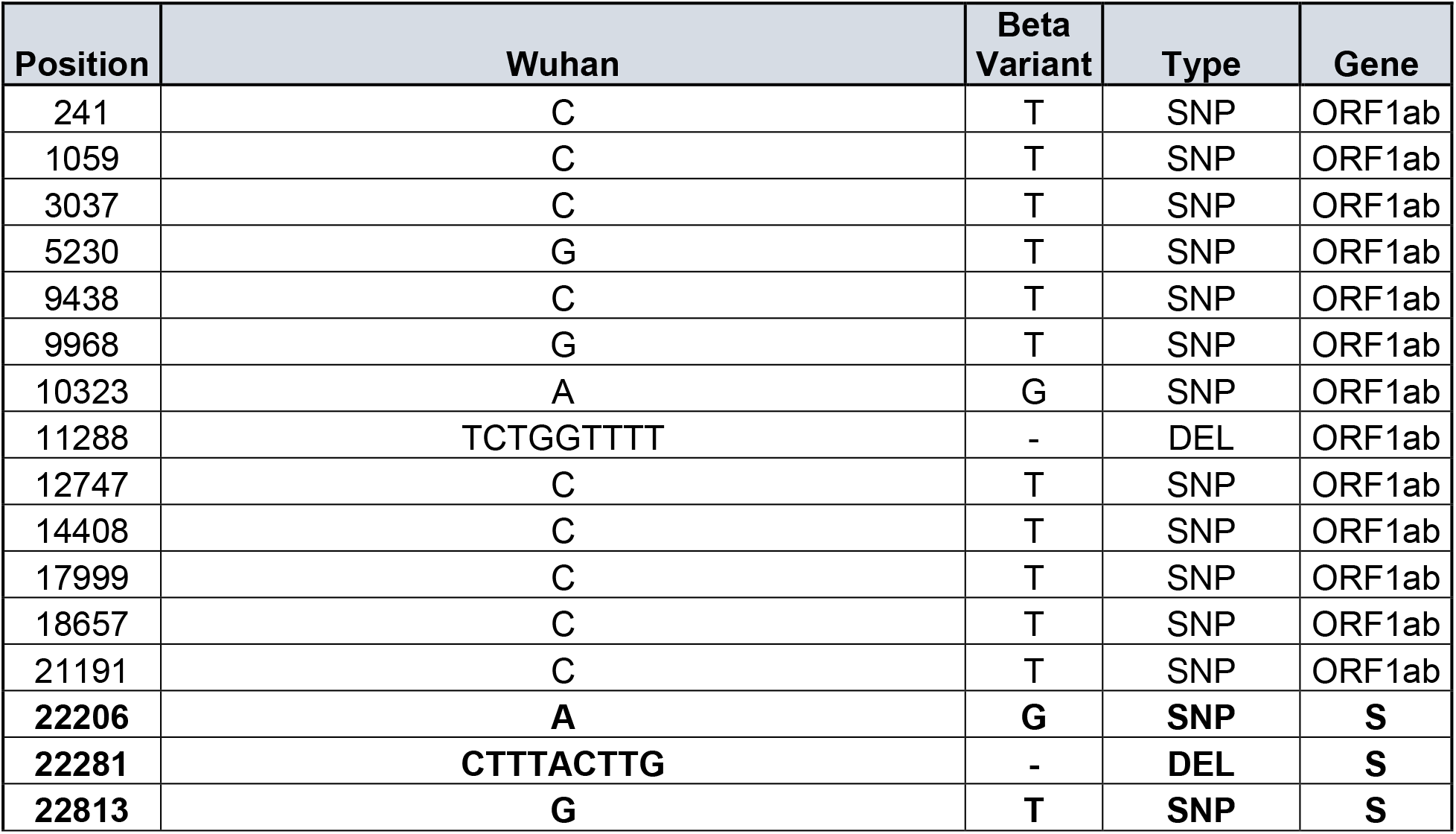
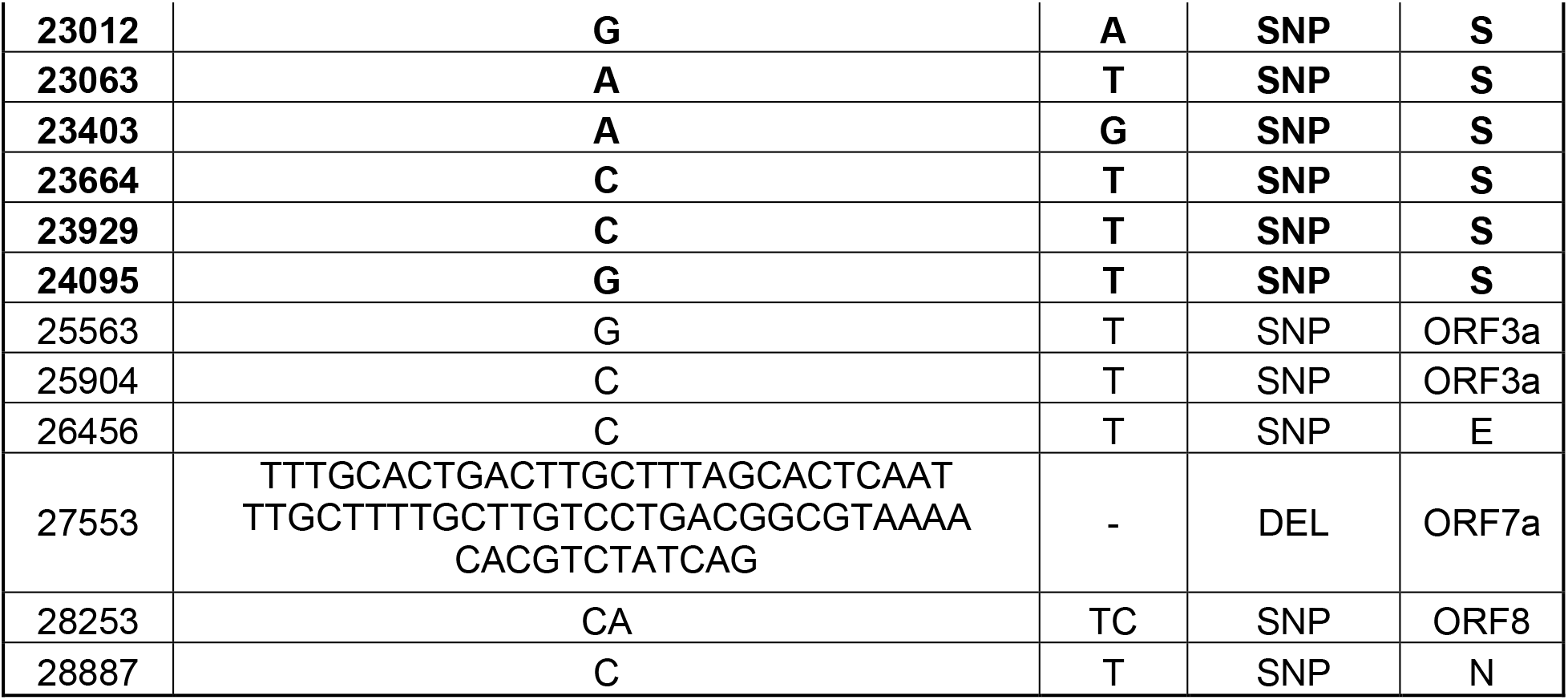

### Preparation of recombinant human ACE2

Clinical grade recombinant human ACE2 (amino acids 18-740) was produced by the contract manufacturer Polymun Scientific (Klosterneuburg, Austria) from CHO cells according to GMP guidelines under serum free conditions and formulated as a physiologic aqueous solution, as described previously (Haschke et al, Clin Pharmacokinet, 2013 and Zoufaly et al, 2021, Lancet Respiratory Medicine).

### ELISA experiments

96-well ELISA plates were coated with 100μL of anti-ACE2 coating antibody (2μg/ml diluted in PBS pH 7.4) over night at room temperature. Following coating, plates were washed 3 times with 300μL of washing buffer (PBS + 0.05% Tween-20) and blocked with 300μL of blocking buffer (1% BSA in PBS, pH 7,4) for 1h at room temperature. After blocking, plates were washed five times and 100μL APN01, diluted in blocking buffer to a concentration of 2μg/mL, was applied to the wells and incubated for 1 hour at room temperature. Subsequently, plates were washed five times with washing buffer and 100μL of SARS-CoV-2 Spike RBD-Variants were added to the plates in triplicates. Following incubation for 1 hour at room temperature, plates were washed five times with washing buffer after which 100μL of anti-Histidine detection antibody (diluted 1/500 in blocking buffer) was added for another hour. After five washing steps, 100μL secondary HRP conjugated antibody was added for one hour at room temperature. Subsequently the plates were washed seven times with washing buffer and 100μL substrate (TMB Microwell Peroxidase Substrate, Seramun Diagnostika #S-001-2-TMB, ready-to-use) was added to the wells. The colorimetric reaction was stopped by the addition of 50μL of 1M sulfuric acid and subsequently analyzed for absorption at 450nm. Blank values were subtracted from data.

### Visualizations of RBDs and full-length Spike protein

Visualizations were rendered with pymol software (the PyMOL Molecular Graphics System, Version 2.4 Schrödinger, LLC), based on a model of the fully glycosylated Spike-ACE2 complex described in Capraz et al., (submitted for publication) and https://covid.molssi.org//models/#spike-protein-in-complex-with-human-ace2-ace2-spike-binding.

### Primers and Antibodies

The following tables lists the primers and Antibodies used in this study:

**Primers.**
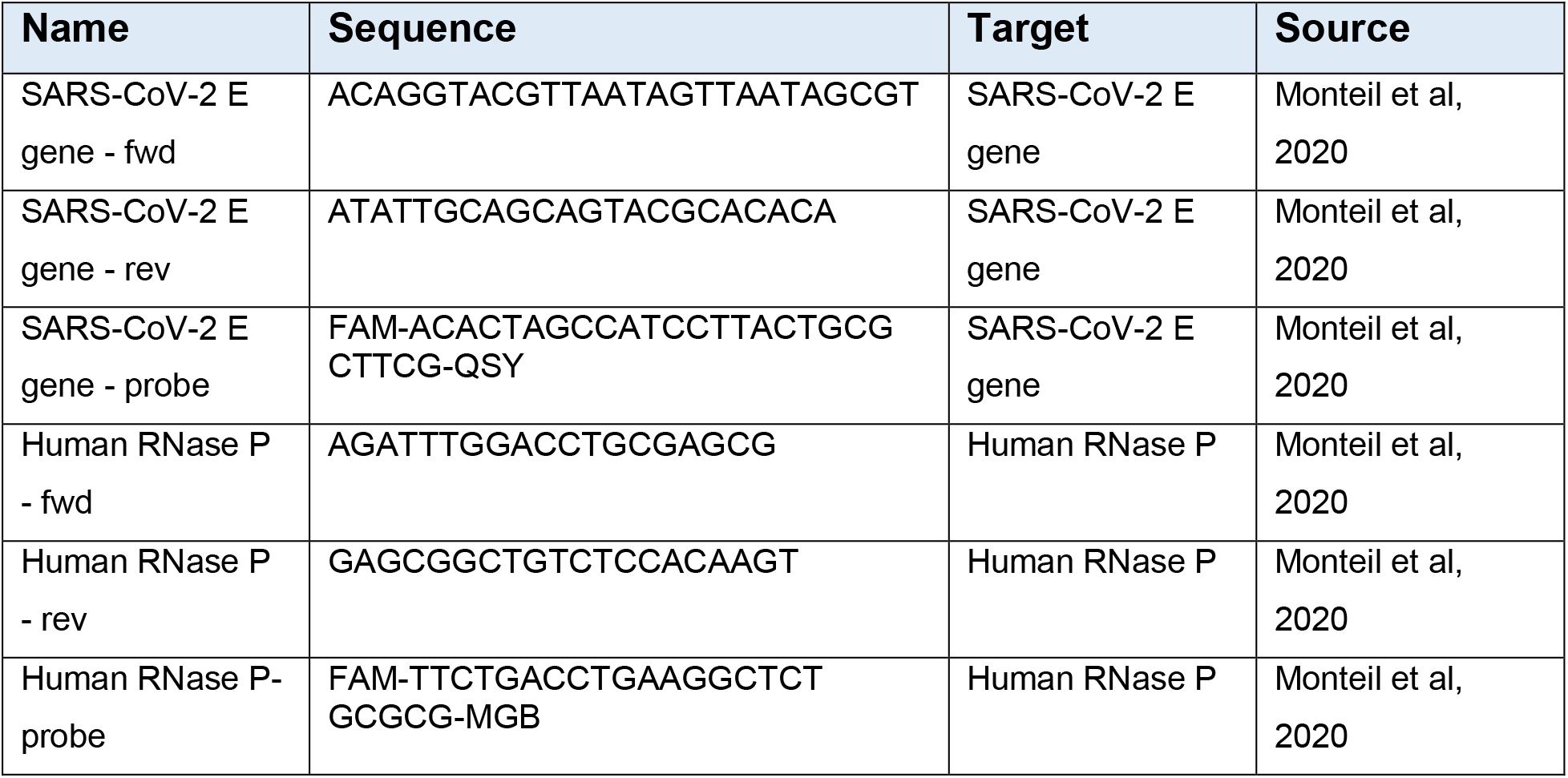

**Antibodies.**

| Name | Vendor | Conjugate | Kit or product number |
| --- | --- | --- | --- |
| ACE2 capture antibody | R&D Systems | unconjugated | DuoSet ELISA Kit, Human ACE2, DY933-05 |
| Mouse anti-His-tag | Biorad | unconjugated | MCA1396 |
| Anti-mouse IgG | CellSignaling | HRP | #7076S |
| SARS-CoV-1 NP<br>antibody (rabbit) | Sino<br>Biological | unconjugated | # 40143-R001 – cross reactive to<br>SARS-CoV-2 |
| Goat anti-rabbit IgG | Life<br>Technologies | Alexa 647 | #A21245 |

